# Scabrous is distributed via signaling filopodia to modulate Notch response during bristle patterning in Drosophila

**DOI:** 10.1101/2023.04.20.537357

**Authors:** Adam Presser, Olivia Freund, Theodora Hassapelis, Ginger L Hunter

## Abstract

During development, tissues must be patterned correctly in order to support tissue function and shape. The sensory bristles of the peripheral nervous system on the thorax of Drosophila melanogaster self-organize from an unpatterned epithelial tissue to a regular spot pattern during pupal stages. Wild type patterning requires Notch-mediated lateral inhibition. Scabrous is a protein that can bind to and modify Notch receptor activity. Scabrous can be secreted, but it is also known to be localized to basal signaling filopodia, or cytonemes, that play a role in long-range Notch signaling. Consistent with previous work, we show that Scabrous is primarily distributed basally, within the range of signaling filopodia extension. We show that signaling filopodia are required for the distribution of Scabrous protein during sensory bristle patterning stages. We show that the Notch response of epithelial cells is sensitive to the level of Scabrous protein being expressed by the sensory bristle precursor cell. Our findings at the cell-level suggest a model for how epithelial cells engaged in lateral inhibition at a distance are sensitive local levels of Scabrous protein.

## Introduction

The timely and ordered patterning of tissues during development contributes to overall tissue homeostasis and animal behavior. The array of microchaete sensory bristles on the dorsal thorax of the fruit fly Drosophila melanogaster is a well-studied example of a self-organizing, repeating spot pattern driven by Notch-mediated lateral inhibition (Furman and Bukharina, 2008; Simpson, 1990; Troost et al., 2015). The thoracic epithelia is initially unpatterned, and most cells are able to commit to either an epithelial or sensory bristle precursor cell fate. Over several hours, bristle precursor cells are specified by low levels of activated Notch and increased expression of proneural genes. Bristle precursor cells engage in lateral inhibition with neighboring epithelial cells. A feature of this system pattern refinement, where cells in the patterning tissue are able to flexibly adjust their cell fate during patterning in order to ensure the correct spacing between bristle precursors (Cohen et al., 2010; Koto et al., 2011). For example, if a bristle precursor cell is ablated, a nearby uncommitted epithelial cell can take its place (Cohen et al., 2010; Hunter et al., 2016). The ability to switch fates is a property of most epithelial cells in the notum and is linked to Notch signaling status and G2-exit. The flexibility of cells fates in the tissue helps to ensure a robust sensory organ pattern. Failure to correctly organize the thoracic sensory bristles can lead to changes in fly behavioral responses (Lacoste et al., 2022). Furthermore the spatial distribution of sensory bristles is under selective pressure as Drosophila species exhibit a variety of bristle densities and organizations (Simpson et al., 1999). Therefore it is important to understand the mechanisms that result in successful tissue patterning.

Theoretical and experimental data demonstrate that molecular modifiers of Notch signaling and cell morphology play roles in the density of the final bristle pattern.

Models clearly support a characteristic length for the lateral inhibition signal beyond adjacent cells, which could be achieved by active cell processes and by diffusible signals (Cohen et al., 2010; Corson et al., 2017; Hadjivasiliou et al., 2016; Hadjivasiliou and Hunter, 2022). Dynamic actin-based signaling filopodia, or cytonemes, have been shown to play a role in the Notch response of epithelial cells during patterning stages (Cohen et al., 2010; Georgiou and Baum, 2010; Hunter et al., 2019). Filopodia structures extend from the basal surface of all notum epithelial cells and permit cells that are 2-3 cell diameters away from each other to contact and therefore engage in Notch signaling. Both Notch receptor and Delta ligand have been shown to localize along the length of signaling filopodia, similar to cytoneme-mediated Notch signaling in other tissues (Boukhatmi et al., 2020; de Joussineau et al., 2003; Huang and Kornberg, 2015; Renaud and Simpson, 2001). Direct evidence of Notch activation along signaling filopodia has not been shown. However, there is genetic evidence that perturbation of actin regulators, and therefore signaling filopodia formation and activity, leads to defects in bristle patterning consistent with mathematical models (Cohen et al., 2010; Georgiou and Baum, 2010; Hunter et al., 2019).

An alternative to active cell processes extending the range of Notch signaling would be the presence of a diffusible regulator of Notch signaling. Scabrous is a fibrinogen-like protein that binds to Notch receptors and plays a role in the formation of a well-spaced array of sensory bristles in the dorsal thorax (Powell et al., 2001; Renaud and Simpson, 2001). Loss of Scabrous expression leads to an increase in bristle density, although how this is achieved is unclear. Scabrous can contribute to the stability of Notch receptor on the cell surface (Powell et al., 2001), however, it can also block Delta-Notch interactions (Lee et al., 2000). There is evidence that Scabrous is a secreted and diffusible Notch regulator, expressed primarily by Notch-inactive, Delta-expressing bristle precursor cells (Baker et al., 1990; Hu et al., 1995; Lee et al., 1996). Other studies indicate that Scabrous is carried via long, actin-rich signaling filopodia (Chou and Chien, 2002; Lacoste et al., 2022), the activity of which are also implicated in establishing bristle density as discussed above (Cohen et al., 2010; Hunter et al., 2019). Signaling filopodia formation and dynamics are Scabrous independent (Renaud and Simpson, 2001) but both are up-regulated in cells that express high levels of Delta ligand (Buffin and Gho, 2010; de Joussineau et al., 2003).

Here we investigate the role of Scabrous in long-range Notch signaling during thoracic bristle patterning. Using genetic and pharmacological strategies, we have found that Scabrous protein is primarily localized to the basal surface of the patterning epithelium, where its distribution is signaling filopodia dependent. We further show that loss of the Scabrous gradient leads to disruption of the Notch response in epithelial cells. We propose that Scabrous helps to coordinate pattern refinement by modifying Notch signaling response in epithelial cells that are distant from Delta-expressing bristle precursor cells.

## Materials and methods

### Fly husbandry

Drosophila stocks were maintained on standard Drosophila food (JazzMix, Fisher Scientific), at 18°C with 24 hour light cycle. Crosses are maintained at room temperature (23-25°C). White pre-pupae were screened against balancers and then aged at 18°C for 24 hours in humidified chambers. Pupae aged to 12-14 hours after pupariation were then dissected for live or fixed imaging protocols.

### Immunofluorescence

Dissected nota of 12-14 hAP pupae were fixed for 20 minutes in 4% paraformaldehyde/1X PBS solution. Tissues were then blocked in 1:1 blocking buffer (5% w/v BSA, 3% FBS in 1X PBS) in 1X PBST at room temperature for 1 hour.

Tissues were then incubated in primary antibody with 5% blocking buffer in 1X PBST for 2 hours at room temperature or 4°C overnight. Tissues were washed twice in 1X PBST for 10 minutes each at room temperature, followed by incubation with secondary antibody in 1X PBST containing fluorescently-labeled phalloidin and DAPI, for 2 hours at room temperature or 4°C overnight. Tissues were washed twice in 1X PBST for 10 minutes each, then equilibrated in 50% glycerol overnight. Nota were then mounted on coverslips and sealed with nail varnish for storage at 4°C until imaged.

### Pharmacological treatments

Cytochalasin B stock solutions were made with DMSO and aliquots maintained at -20°C. For experimental treatments, Cytochalasin was diluted to 20ug/mL in room temperature modified Clone 8 medium (2.5% Insect media supplement, 2% FBS, in 1X Schneider’s Medium)(Loubéry and González-Gaitán, 2014). The same volume of DMSO was added to medium for control treatments. The genotype used for this experiment is neur-GAL4, UAS-GMCA/+. Dissected nota were incubated at room temperature, in a humidified chamber, with either 20ug/mL cytochalasin B or control solution for the time points described in Figure 2. At the end of the incubation period, the solution was removed and fixative was added, followed by immunofluorescence protocols described above. For live imaging, fully dissected nota were held in place on a Matek dish by the fibrin clot method (Coumailleau et al., 2009) in a small volume of modified Clone 8 medium. An equal volume of medium containing vehicle or cytochalasin was added, to reach the appropriate final concentration.

### Imaging

Samples were imaged on a either a Lieca DMi8 SPE confocal microscope using LASX software, or Nikon C2+ confocal microscope using NIS Elements. The pupal cases of live 12-14 hAP pupae were removed, exposing the head and thorax. A coverslip coated with a thin film of Halocarbon 27 oil (Sigma) was placed on top of spacers such that only the dorsal thorax contacted the coverslip (as previously published (Loubéry and González-Gaitán, 2014)). Live pupae were imaged using a x40 (0.8 NA) air objective. Fixed tissues were imaged using a x40 (1.15NA), x63 (1.3 NA) oil objectives or Nikon C2+ using a x40 (1.3 NA). Ex vivo pharmacological experiments were performed at room temperature on a Nikon EclipseTi, x60 (1.4 NA).

### Quantification and statistical analysis

All image analysis was performed in FIJI/ImageJ. Figure 1B: apical and basal distribution of Scabrous was determined by analyzing z-slices apical or basal to the cell nucleus marked by anti-senseless. Figure 1C: Area adjusted number of Scabrous puncta was calculated by the equation f(x) = x / ((d+1)2-d2). Figure 2C-D: A circular ROI with radius of 15 μm was centered on a bristle precursor labeled with anti-GFP. The number of TRITC +, GFP-puncta within this ROI was counted as extracellular Scabrous and TRITC+, GFP+ puncta were counted as intracellular Scabrous.

**Fig 1.**
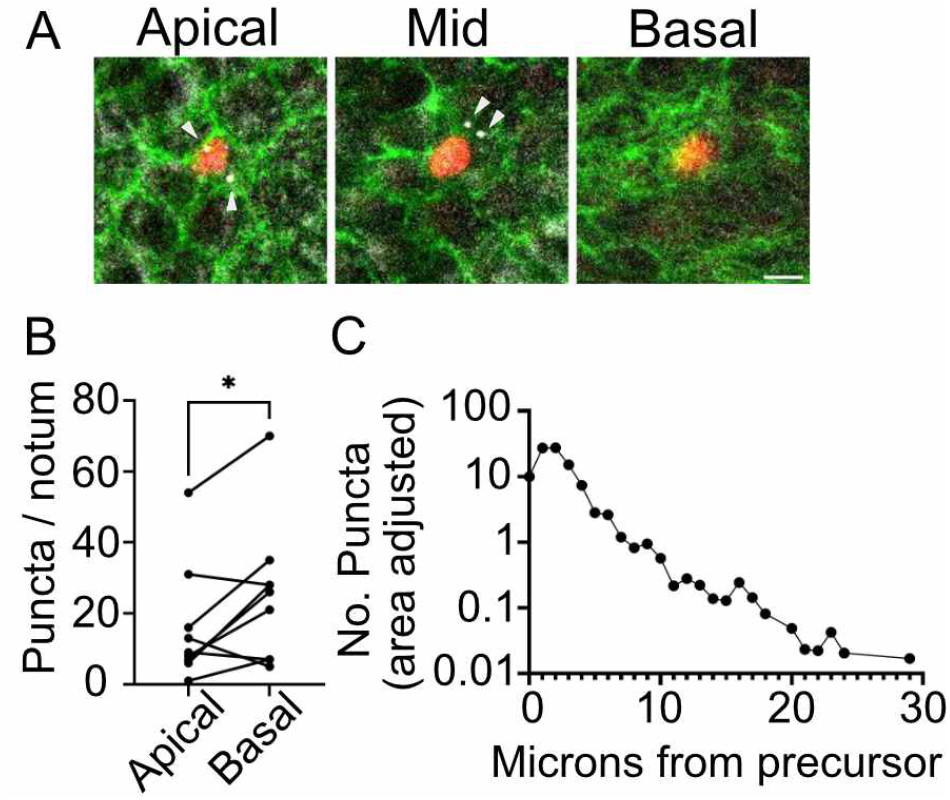
Spatial distribution of Scabrous protein during bristle patterning stages. (A) Example of apicobasal distribution of Scabrous protein (arrowheads, white puncta, anti-Scabrous) at a bristle precursor cell, marked by red nuclei (anti-senseless). Phalloidin is used to stain filamentous actin. Scale bar, 5 μm. (B) Pairwise quantification of Scabrous puncta at apical or basal z-planes for n = 9 nota. *, p = 0.04, r = 0.83 by paired student’s t-test. (C) XY distribution of Scabrous puncta relative to the center of bristle precursor cells (0 μm). The number of puncta are area adjusted, across n = 9 nota.

Maximum intensity projections were used for this analysis. Figure 3B: An ROI with a height and width of 3 μm was centered on the nuclei of NsfGFP expressing epithelial cells, and the average fluorescence intensity was measured. Adjacent or distant cell nuclei were determined by relative positioning to the nearest RFP+ nuclei (bristle cells are marked by neur-H2B:mRFP). Fluorescence intensity measurements were taken from a single z-slices at the middle of cells, due to issues with Scabrous-GFP overexpression obstructing nuclear NsfGFP measurements in maximum intensity projections.

Statistical analysis was performed using GraphPad Prism. Specific statistical tests are described in the figure legends.

### Drosophila stocks

NsfGFP, neur:H2B-RFP/CyOGFP; pannier-GAL4/TM6BTb (Hunter et al., 2016) UAS-scabrous RNAi / CyOGFP (BDSC). UAS-scabrous GFP / CyOGFP (BDSC). UAS-white RNAi (BDSC). Neur-GAL4, UAS-GMCA /TM6BTb (Georgiou and Baum, 2010). W1118 (BDSC).

### Antibodies and stains

Mouse anti-scabrous 1:500, DSHB (Lee et al., 1996). Guinea pig anti-senseless 1:2000, Bellen Lab (Nolo et al., 2000). Chicken anti-GFP 1:1000, EMD Millipore AB16901. AF488 anti-chicken 1:2000, Jackson Immunological 703-545-155. 555 anti-guinea pig 1:2000, Invitrogen A21435. TRITC anti-mouse 1:2000, Jackson Immunological 715-025-151. 647 anti-mouse 1:2000, Invitrovgen A48289. Phalloidin (555 or 670) 1:500, Cytoskeleton Inc PHDH1, PHDN1. DAPI 1:1000, Thermoscientific 62248

### Pharmacological reagents

Cytochalasin B 20ug/mL (Tocris 5474). DMSO (Sigma D5879)

## Results and Discussion

### Scabrous distribution in vivo

In order to determine the contribution of Scabrous to long-range Notch signaling, we first determined the spatial distribution of endogenous Scabrous protein in the patterning notum (Figure 1). We stained for endogenous Scabrous protein in nota at bristle patterning stages (14 hours after pupariation, hAP). Endogenously expressed Scabrous protein is primarily localized to aggregates in bristle precursor cells (arrowheads, Figure 1A), which likely represents localization to vesicles, as previously described (Renaud and Simpson, 2001). We next determined the apical-basal distribution of Scabrous protein. We used a fluorescent phalloidin F-actin stain to visualize the apical junctions as well as the basal signaling filopodia (Figure 1A). We find that Scabrous puncta distribution is skewed towards the basal side of the tissue (Figure 1B). We then determined the overall distribution of Scabrous puncta relative to the nearest bristle precursor cell nuclei. Scabrous puncta can be observed as far as 2-3 cell diameters away (20-30 microns, Figure 1C), although most are localized within 10 microns of the bristle cell nuclei. Together these observations are consistent with previous reports that Scabrous-GFP can localize to the basal signaling filopodia of bristle precursor cells (Lacoste et al., 2022), and that Scabrous acts cell non-autonomously (Renaud and Simpson, 2001). Interestingly, the average length of signaling filopodia extending from the basal surface of bristle precursor cells is 10 μm (Hunter et al., 2019). Therefore, the XY distribution of Scabrous could be consistent with either active or passive processes.

### Scabrous distribution is dependent on the activity of signaling filopodia

We and others have observed that Scabrous is distributed both apically and basally in the patterning notum. In a previous study, Scabrous was shown to localize along the length of basal signaling filopodia in the patterning notum (Lacoste et al., 2022). It is currently unknown if Scabrous is distributed via these signaling filopodia, or through a diffusion-based process. Importantly, while both Scabrous and signaling filopodia are up-regulated in cells that express Delta, the formation of signaling filopodia is not Scabrous dependent (de Joussineau et al., 2003; Renaud and Simpson, 2001). Therefore we next asked whether the distribution of Scabrous in vivo is dependent on the activity of signaling filopodia.

Since signaling filopodia are actin-dependent structures, we used the filamentous actin (F-actin) inhibitor Cytochalasin B to acutely disrupt actin during patterning stages (MacLean-Fletcher and Pollard, 1980). Previously, we had adapted an ex vivo technique for the pharmacological perturbation of nota (Hunter et al., 2019; Loubéry and González-Gaitán, 2014). We found that treatment with Cytochalasin B leads to the rapid disassembly of signaling filopodia in notum explants, compared to DMSO controls (Figure 2A). F-actin disruption, visualized by loss of LifeAct-Ruby containing cell protrusions, was apparent by ¡ 60 seconds after addition of Cytochalasin B. We hypothesized that if signaling filopodia are essential to Scabrous distribution, then loss of signaling filopodia would lead to an decrease in the number of Scabrous puncta at a distance from bristle precursor cells. In order to determine when and how the distribution of Scabrous changed in response to F-actin disruption, we exposed dissected nota to Cytochalasin B or DMSO (Figure 2B-D) at designated time-points. We find that long-term incubation with DMSO alone do not disrupt the formation of F-actin based signaling filopodia (Figure 2B). For these experiments, we used the neur-GAL4, UAS-GMCA (Cohen et al., 2010; Georgiou and Baum, 2010; Hunter et al., 2019) genetic background that labels bristle precursor cells only with a GFP-tagged actin binding domain of Moesin (GMCA;(Edwards et al., 1997)).

**Fig 2.**
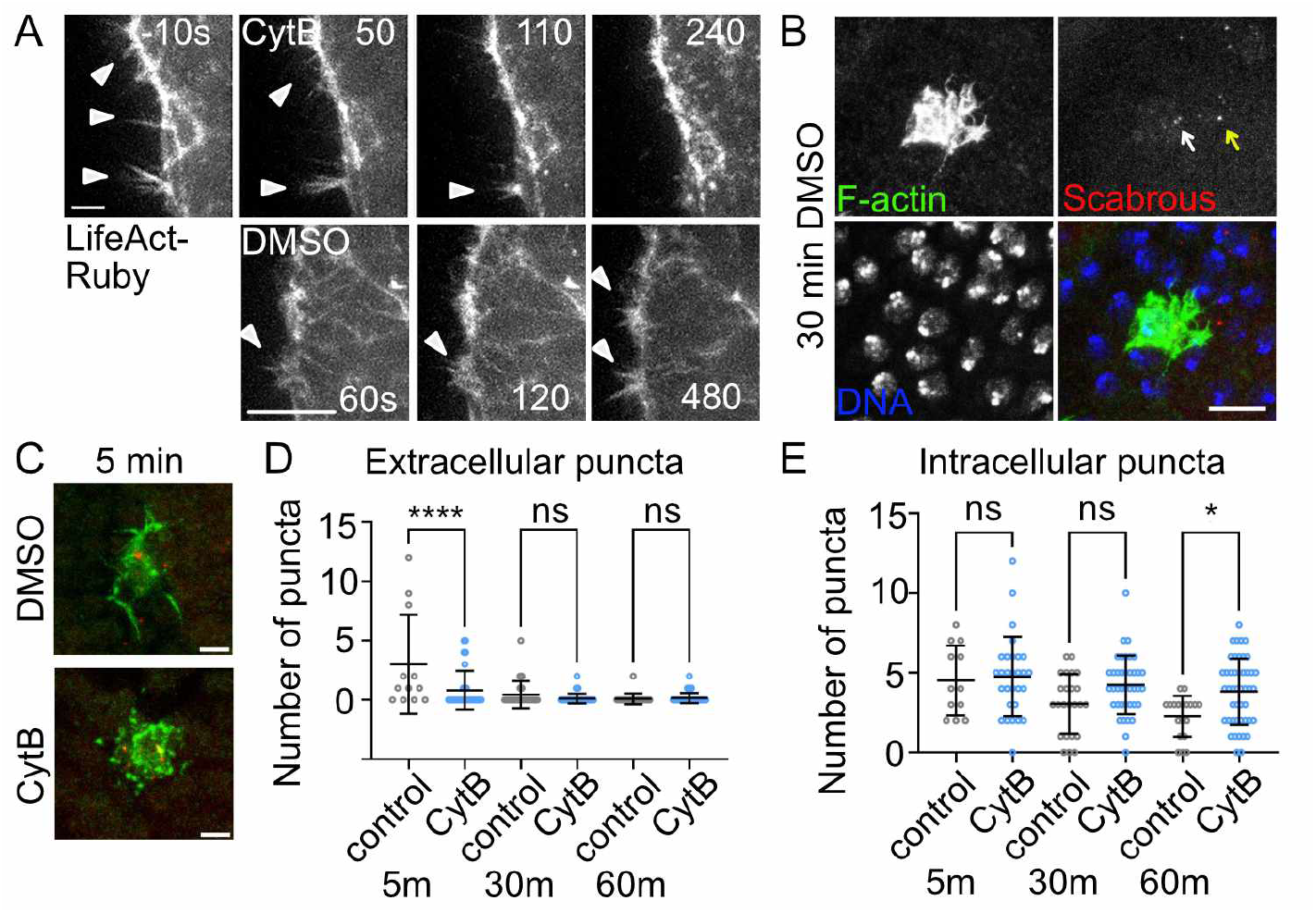
Signaling filopodia are required for the extracellular distribution of Scabrous. (A) Timelapse panels of nota explants expressing the filamentous actin marker LifeAct-Ruby and treated with either DMSO or Cytochalasin B. Time in seconds. Either treatment was added at time = 0 seconds. Arrowheads point to individual signaling filopodia on the basal surface. Scale bars, 4 μm (CytB) and 10 μm (DMSO). (B) Immunofluorescence image of a sensory bristle precursor cell from a nota under control conditions for 30 minutes prior to fixation. Signaling filopodia are still visible in F-actin panel. White arrow in Scabrous panel points to intracellular Scabrous. Yellow arrow in Scabrous panel points to extracellular Scabrous. Scale bar, 10 μm. (C) Example cell morphologies upon DMSO control or Cytochalasin B treatments for 5 minutes. Green, anti-GFP (GFP-actin binding domain of moesin labeling F-actin). Red, anti-Scabrous. Scale bar, 5 μm. (D) Quantification of extracellular Scabrous puncta within a 15 μm radius of a bristle precursor cell in nota treated with either DMSO for 5 (n = 12 cells), 30 (n = 19 cells), or 60 (n = 39 cells) minutes, or cytochalasin B for 5 (n = 29 cells), 30 (n = 18 cells), or 60 (n = 46 cells) minutes prior to fixation. A minimum of 5 nota were dissected and treated for each time point and condition. ****, p¡0.0001 by ordinary one-way ANOVA with correction for multiple pairwise comparisons. (E) Quantification of intracellular Scabrous puncta for the same cells in (C). *, p = 0.0162 by ordinary one-way ANOVA with correction for multiple pairwise comparisons. Ns = not significant.

Treatment of dissected nota with Cytochalasin B leads to disruption of the F-actin cytoskeleton by the first time point observed (5 minutes), indicated by disruption of GMCA localization in bristle precursor cells (Figure 2C). To quantify Scabrous distribution under disrupted or control conditions, we count the number of Scabrous-positive puncta within signaling filopodia range of a bristle precursor cells. The average maximum length of signaling filopodia extending from bristle precursor cells is 10 μm (Cohen et al., 2010; Hunter et al., 2019), and the average diameter of a bristle precursor cell prior to cell division is 10 μm. Therefore we performed our analysis within a 15 μm radius of each bristle precursor cell. Scabrous-positive puncta that did not co-localize with anti-GFP were considered extracellular to the bristle precursor cells, and Scabrous-positive puncta that co-localized with anti-GFP were considered intracellular (Figure 2B, arrows). We found that the number of extracellular Scabrous puncta was decreased within 5 minutes of cytochalasin B treatment compared to controls (Figure 2D). At longer term treatments, this effect decreased. This finding supports a role for signaling filopodia in the distribution of Scabrous. It remains to be seen if the activity of Scabrous on Notch signaling requires it to be transferred from the signaling to the receiving cell, similar to what has been observed for Sonic Hedgehog signaling along cytonemes (Hall et al., 2021).

Several steps in the secretory pathway and exocytosis require the activity of actin cytoskeleton (Chakrabarti et al., 2021; Sokac and Bement, 2006). We would expect that treatment of tissues with cytochalasin B would also potentially disrupt the secretion of diffusible Scabrous (Giagtzoglou et al., 2013). We therefore asked if cytochalasin B treatments led to differences in intracellular Scabrous puncta. We predicted we would see increased intracellular localization of Scabrous if its secretion were blocked by actin disruption. At 5 minutes of cytochalasin B treatment, we observe no differences in the number of intracellular Scabrous puncta relative to controls (Figure 2E). We observed increased levels of intracellular Scabrous puncta when tissues are treated with cytochalasin B for longer periods of time (60 minutes, Figure 2E). This result indicates that disruption of the actin cytoskeleton is having an effect on the retention of Scabrous in the bristle precursor cell, at longer timescales. Altogether, our data suggest that the extracellular distribution of Scabrous at shorter timescales requires the activity of signaling filopodia.

### Changes in Scabrous expression levels leads to changes in Notch signaling

Scabrous is known to directly bind to and modulate Notch signaling (Mok et al., 2005; Powell et al., 2001), although its effect on Notch response is unclear. Expression of Scabrous in wing or eye disc clones leads to inactivation of Notch signaling (Lee et al., 2000; Powell et al., 2001). In the notum, the Scabrous phenotype resembles the Notch or Delta hypomorph phenotypes of increased bristles (Renaud and Simpson, 2001), suggesting that it positively regulates Notch activation in the notum. A key difference between these phenotypes is bristle spacing: adjacent bristles are not observed in Scabrous mutant tissues, whereas they are common in Delta or Notch mutant tissues (Heitzler and Simpson, 1991; Renaud and Simpson, 2001). Scabrous could exclusively play a role in the Notch activity of epithelial cells distant from the bristle precursor.

Alternatively, there could be mechanisms that make epithelial cells adjacent to the bristle precursor less sensitive to Scabrous activity. Based on these studies, we hypothesized that decreased Scabrous expression would lead to decreased Notch response in cells that are distant from the source of Delta signal. This could account for the increased number of Notch-inactive cells, and increase in bristle density, in Scabrous mutant or knockdown tissues. Furthermore, if Scabrous protein is distributed by signaling filopodia, we expect to see a strong effect in distant epithelial cells.

To test our hypothesis, we modulated the level of Scabrous expression in nota that expressed a transcriptional reporter of Notch signaling, NsfGFP (Hunter et al., 2016). Here, nuclear GFP is expressed downstream of a minimal Notch responsive promoter, such that increased levels of nuclear GFP fluorescence is associated with higher Notch signaling activity. We used RNAi to decrease the level of Scabrous throughout the notum. We then quantified the nuclear NsfGFP levels in two populations of epithelial cells: (1) adjacent cells, which are those that share a large cell-cell interface with the Delta-expressing bristle precursor cell; (2) distant cells, which are those that are at least one nuclei removed from the bristle precursor cell. Cells more than one nuclei removed from a bristle precursor cell are rare in these modified tissues, as the phenotype of over-or under-expressing Scabrous is to increase the density of bristle precursor cells. Both adjacent and distant epithelial cells are capable of contacting nearby bristle precursor cells via signaling filopodia. However, distant epithelial cells may only use signaling filopodia to engage in contact-mediated Notch signaling with the bristle precursor cells. We measured the nuclear NsfGFP fluorescence levels in 14 hAP nota expressing UAS-scabrous RNAi to decrease Scabrous expression, compared to UAS-white RNAi expressing control, both under the pannier-GAL4 driver which expresses throughout the central notum (Figure 3A).

**Fig 3.**
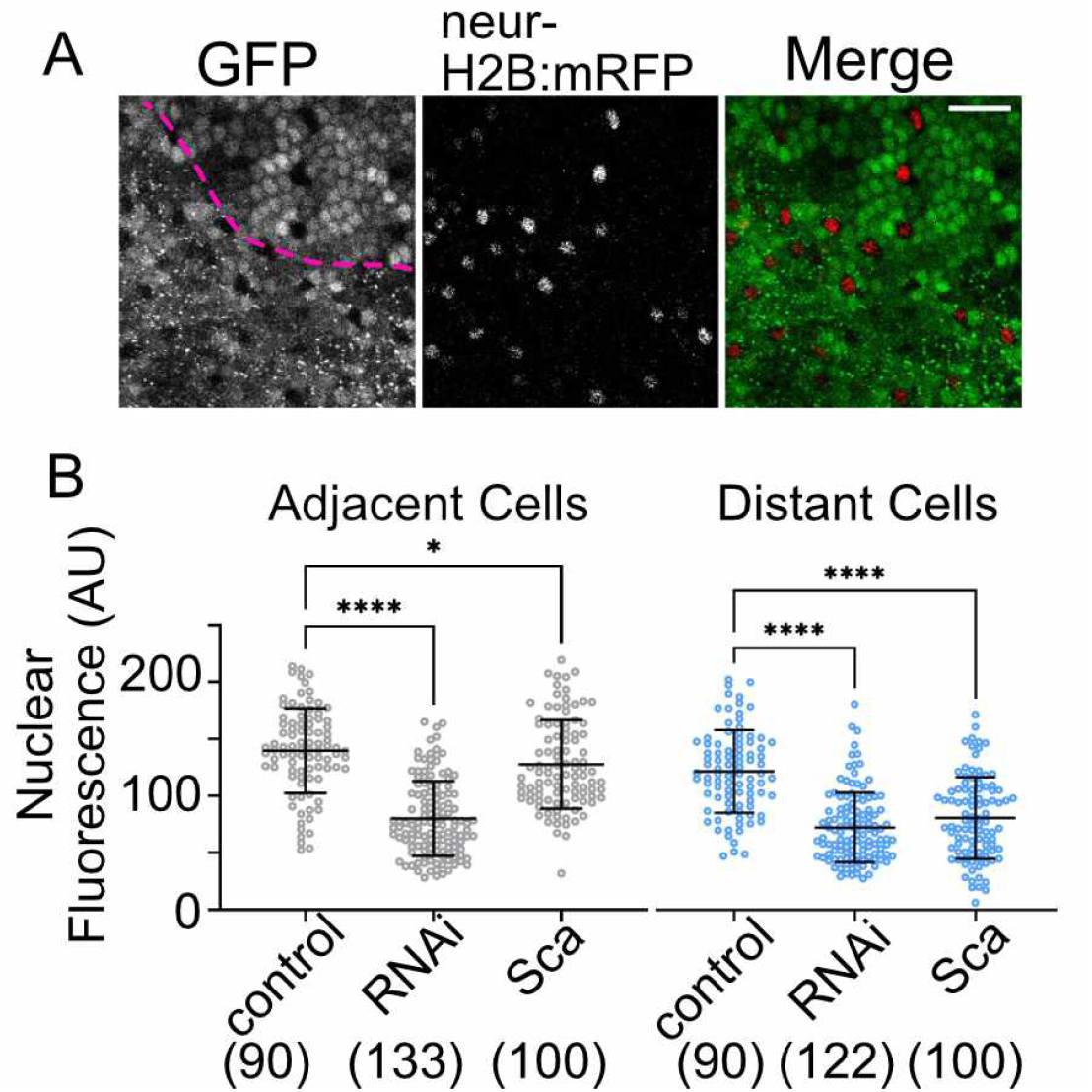
Local Scabrous expression levels are essential for robust Notch response. (A) Example of a live pupal notum expressing UAS-Scabrous GFP in the pannier-GAL4 domain. Lateral edge of the pannier domain is marked by the dashed pink line. All Notch activated cells express the NsfGFP transcriptional reporter, which localizes GFP signal to the nucleus. Bristle precursor cells do no express NsfGFP but do express the neuralized reporter, neur-H2B:mRFP, marking their nuclei. Scale bar, 25 μm. (B) Nuclear GFP fluorescence was measured for cells adjacent to a bristle precursor cell or distant to a bristle precursor cell (+1 nuclei removed) in 14 hAP nota expressing NsfGFP. Tissues were either expressing lower amounts of Scabrous (RNAi = pannier-GAL4 ¿ UAS-Scabrous RNAi), control levels of Scabrous (control = pannier GAL4 ¿ UAS-white RNAi), or elevated levels of Scabrous (Sca = pannier GAL4 ¿ UAS-Scabrous GFP). ****, p¡0.0001 and *, p=0.04 by ordinary one-way ANOVA with multiple comparisons. (n) = number of nuclei measured. A minimum of 3 animals were imaged for each genotype.

In control tissues, epithelial cells adjacent to bristle precursor cells have a higher level of nuclear NsfGFP reporter than distant cells (Hunter et al., 2016). This has been interpreted to be due to increased surface contact area between adjacent epithelial cells and bristle precursor cells leading to increased Notch-Delta engagement and activation (Shaya et al., 2017). When Scabrous levels are decreased via the expression of Scabrous RNAi, we observe that nuclear NsfGFP reporter levels decrease in both adjacent and distant epithelial cells (Figure 3B, RNAi). This suggests that Scabrous acts as a positive regulator of Notch activity in the notum. In the context of previously published results showing that Scabrous mutants have decreased spacing between bristles, but never adjacent bristles, suggests either that (1) the level of Notch activation in adjacent epithelial cell neighbors surpasses a threshold for acquiring the epithelial cell fate or (2) a low, but sufficient, level of Scabrous still being expressed and promoting Notch activation. Due to the short predicted half-life of Scabrous protein (Ellis et al., 1994) as well as the ability of pnr-GAL4¿ UAS-Scabrous RNAi nota to display a Scabrous mutant phenotype of increased bristle density, we favor the threshold model.

Previously it was shown that over-expression of Scabrous throughout a patterning tissue is unable to rescue spacing phenotypes (Ellis et al., 1994; Renaud and Simpson, 2001). It was proposed that this is because local differences in Scabrous levels play a key role in patterning and lateral inhibition. We found that when Scabrous was over-expressed throughout the notum, nuclear NsfGFP reporter levels were decreased relative to controls (Figure 3B, Sca). Adjacent epithelial cells in Scabrous over-expressing tissues exhibit a Notch activity that is only slightly decreased relative to controls. Distant epithelial cells in Scabrous over-expressing tissues exhibit strong decreases in Notch activity relative to controls, comparable to distant cells in Scabrous RNAi expressing tissues. Our results are similar to those observed in the wing disc (Lee et al., 2000), where ectopic expression of Scabrous protein led to the down-regulation of genes downstream of Notch activation. Our results give cell level detail for earlier observations that loss of Scabrous expression and uniform Scabrous over-expression both lead to the same spacing phenotype (Ellis et al., 1994; Renaud and Simpson, 2001).

The decreased Notch response in distant epithelial cells for both RNAi and over-expression conditions may be sufficiently low as to allow cell fate switching, where these distant epithelial cells become bristle precursors.

It remains unclear as to why epithelial cells adjacent to bristle precursors are less affected by over-expression of Scabrous compared to epithelial cells distant to bristle precursors. A major difference between the two epithelial cell types is surface area of contact with the nearest Delta-expressing bristle precursor cell. Assuming both epithelial cell types express Notch receptor at similar levels, contact area differences may have significant impact on the relative numbers of free or engaged Notch receptors. Notch receptors on the cell surface may engage with ligand to be activated in trans-, be inhibited in cis-(Sprinzak et al., 2010), or be endocytosed and turned over without engaging Delta ligand (Sakata et al., 2004). Previous studies show that Scabrous stabilizes cell surface Notch, and that the binding of Notch with Scabrous precludes binding of Notch with Delta ligand (Lee et al., 2000; Powell et al., 2001). Together with the findings in our study, we propose a model in which (1) loss of Scabrous expression destabilizes cell surface Notch. In combination with Notch receptor turnover, this state leads to decreased engagement with Delta and lower Notch response in epithelial cells (as in Figure 3B RNAi). (2) Over-expression of Scabrous leads to stabilization of Notch on the cell surface but prevents the ligand from binding to the receptor, also leading to lower Notch response in epithelial cells (as in Figure 3B Sca). The observation that adjacent cells are not as strongly affected as distant cells may be related to the relative affinity of Notch for Delta vs Scabrous. If the relative affinity of Notch for Delta is higher than for Scabrous, and since adjacent epithelial cells have more opportunities to interact with Delta ligand on contact surfaces than distant epithelial cells do via signaling filopodia contacts alone, we would expect to see a higher Notch response in adjacent cells.

## Conclusion

In summary, our evidence demonstrates that Scabrous can be distributed basally via signaling filopodia during the patterning of the notum epithelium. Consistent with previous research, Scabrous is important for enhancing Notch signaling during lateral inhibition. Our analysis at the level of individual cells clarifies earlier observations that decreasing and increasing levels of Scabrous both result in similar tissue patterning defects. Future work will be needed to address the detailed mechanism by which Scabrous modulates Notch signaling during bristle patterning, and how its signal is transmitted via signaling filopodia.

## Acknowledgments

We thank the laboratory of Dr Hugo Bellen (Baylor College of Medicine) for the Guinea pig anti-scabrous antibody. We thank Dr Jill Pflugheber (St. Lawrence University) for emergency help with microscopy. We thank Dr Edward Giniger (NINDS, NIH) for postdoctoral training support and support for the ex vivo imaging experiments. We thank the Hunter laboratory for their review and comments on this manuscript. Stocks obtained from the Bloomington Drosophila Stock Center (NIH P40OD018537) were used in this study. The anti-Scabrous monoclonal antibody was obtained from the Developmental Studies Hybridoma Bank, created by the NICHD of the NIH and maintained at The University of Iowa, Department of Biology, Iowa City, IA 52242.

## Notes

### Competing Interest Statement

The authors have declared no competing interest.

## References

1. Baker, N.E., Mlodzik, M., Rubin, G.M., 1990. Spacing Differentiation in the Developing Drosophila Eye: a Fibrinogen-related Lateral Inhibitor Encoded by scabrous. Science 250, 1370–1377. https://doi.org/10.1126/science.2175046.

2. Boukhatmi, H., Martins, T., Pillidge, Z., Kamenova, T., Bray, S., 2020. Notch Mediates Inter-tissue Communication to Promote Tumorigenesis. Curr. Biol. 30, 1809–1820.e4. https://doi.org/10.1016/j.cub.2020.02.088

3. Buffin, E., Gho, M., 2010.Laser Microdissection of Sensory Organ Precursor Cells of Drosophila Microchaetes. PLoS ONE 5, e9285 https://doi.org/10.1371/journal.pone.0009285

4. Chakrabarti, R., Lee, M., Higgs, H.N., 2021. Multiple roles for actin in secretory and endocytic pathways. Curr. Biol. 31, R603–R618 https://doi.org/10.1016/j.cub.2021.03.038

5. Chou, Y.-H., Chien, C.-T., 2002. Scabrous Controls Ommatidial Rotation in the Drosophila Compound Eye. Dev. Cell 3, 839–850. https://doi.org/10.1016/S1534-5807(02)00362-3

6. Cohen, M., Georgiou, M., Stevenson, N.L., Miodownik, M., Baum, B., 2010. Dynamic Filopodia Transmit Intermittent Delta-Notch Signaling to Drive Pattern Refinement during Lateral Inhibition. Dev. Cell 19, 78–89. https://doi.org/10.1016/j.devcel.2010.06.006

7. Corson, F., Couturier, L., Rouault, H., Mazouni, K., Schweisguth, F., 2017. Self-organized Notch dynamics generate stereotyped sensory organ patterns in Drosophila. Science 356, eaai7407. https://doi.org/10.1126/science.aai7407

8. Coumailleau, F., Fürthauer, M., Knoblich, J.A., González-Gaitán, M., 2009. Directional Delta and Notch trafficking in Sara endosomes during asymmetric cell division. Nature 458, 1051–1055. https://doi.org/10.1038/nature07854

9. de Joussineau, C., Soulé, J., Martin, M., Anguille, C., Montcourrier, P., Alexandre, D., 2003. Delta-promoted filopodia mediate long-range lateral inhibition in Drosophila. Nature 426, 555–559. https://doi.org/10.1038/nature02157

10. Edwards, K.A., Demsky, M., Montague, R.A., Weymouth, N., Kiehart, D.P., 1997. GFP-Moesin Illuminates Actin Cytoskeleton Dynamics in Living Tissue and Demonstrates Cell Shape Changes during Morphogenesis in Drosophila. Dev. Biol. 191, 103–117. https://doi.org/10.1006/dbio.1997.8707

11. Ellis, M.C., Weber, U., Wiersdorff, V., Mlodzik, M., 1994. Confrontation of scabrous expressing and non-expressing cells is essential for normal ommatidial spacing in the Drosophila eye. Development 120, 1959–1969. https://doi.org/10.1242/dev.120.7.1959

12. Furman, D., Bukharina, T., 2008. How Drosophila melanogaster Forms its Mechanoreceptors. Curr. Genomics 9, 312–323. https://doi.org/10.2174/138920208785133271

13. Georgiou, M., Baum, B., 2010. Polarity proteins and Rho GTPases cooperate to spatially organise epithelial actin-based protrusions. J. Cell Sci. 11. DOI: 10.1242/jcs.060772

14. Giagtzoglou, N., Li, T., Yamamoto, S., Bellen, H.J., 2013. dEHBP1 regulates Scabrous secretion during Notch mediated lateral inhibition. J. Cell Sci. jcs.126292. https://doi.org/10.1242/jcs.126292

15. Hadjivasiliou, Z., Hunter, G., 2022. Talking to your neighbors across scales: Long-distance Notch signaling during patterning, in: Current Topics in Developmental Biology. Elsevier, pp. 299–334. https://doi.org/10.1016/bs.ctdb.2022.04.002

16. Hadjivasiliou, Z., Hunter, G.L., Baum, B., 2016. A new mechanism for spatial pattern formation via lateral and protrusion-mediated lateral signalling. J. R. Soc. Interface 13. https://doi.org/10.1098/rsif.2016.0484

17. Hall, E.T., Dillard, M.E., Stewart, D.P., Zhang, Y., Wagner, B., Levine, R.M., Pruett-Miller, S.M., Sykes, A., Temirov, J., Cheney, R.E., Mori, M., Robinson, C.G., Ogden, S.K., 2021. Cytoneme delivery of Sonic Hedgehog from ligand-producing cells requires Myosin 10 and a Dispatched-BOC/CDON co-receptor complex. eLife 10, e61432. https://doi.org/10.7554/eLife.61432

18. Heitzler, P., Simpson, P., 1991. The choice of cell fate in the epidermis of Drosophila. Cell 64, 1083–1092. https://doi.org/10.1016/0092-8674(91)90263-X

19. Hu, X., Lee, E.C., Baker, N.E., 1995. Molecular analysis of scabrous mutant alleles from Drosophila melanogaster indicates a secreted protein with two functional domains. Genetics 141, 607–617. https://doi.org/10.1093/genetics/141.2.607

20. Huang, H., Kornberg, T.B., 2015. Myoblast cytonemes mediate Wg signaling from the wing imaginal disc and Delta-Notch signaling to the air sac primordium. eLife 4. https://doi.org/10.7554/eLife.06114

21. Hunter, G.L., Hadjivasiliou, Z., Bonin, H., He, L., Perrimon, N., Charras, G., Baum, B., 2016. Coordinated control of Notch/Delta signalling and cell cycle progression drives lateral inhibition-mediated tissue patterning. Development 143, 2305–2310. https://doi.org/10.1242/dev.134213

22. Hunter, G.L., He, L., Perrimon, N., Charras, G., Giniger, E., Baum, B., 2019. A role for actomyosin contractility in Notch signaling. BMC Biol. 17. https://doi.org/10.1186/s12915-019-0625-9

23. Koto, A., Kuranaga, E., Miura, M., 2011. Apoptosis Ensures Spacing Pattern Formation of Drosophila Sensory Organs. Curr. Biol. 21, 278–287. https://doi.org/10.1016/j.cub.2011.01.015

24. Lacoste, J., Soula, H., Burg, A., Audibert, A., Darnat, P., Gho, M., Louvet-Vallée, S., 2022. A neural progenitor mitotic wave is required for asynchronous axon outgrowth and morphology. eLife 11, e75746. https://doi.org/10.7554/eLife.75746

25. Lee, E.C., Hu, X., Yu, S.Y., Baker, N.E., 1996. The scabrous gene encodes a secreted glycoprotein dimer and regulates proneural development in Drosophila eyes. Mol. Cell. Biol. 16, 1179–1188. https://doi.org/10.1128/MCB.16.3.1179

26. Lee, E.-C., Yu, S.-Y., Baker, N.E., 2000. The Scabrous protein can act as an extracellular antagonist of Notch signaling in the Drosophila wing. Curr. Biol. 10, 931–S2. https://doi.org/10.1016/S0960-9822(00)00622-9

27. Loubéry, S., González-Gaitán, M., 2014. Monitoring Notch/Delta Endosomal Trafficking and Signaling in Drosophila, in: Methods in Enzymology. Elsevier, pp. 301–321. https://doi.org/10.1016/B978-0-12-397926-1.00017-2

28. MacLean-Fletcher, S., Pollard, 1980. Mechanism of action of cytochalasin B on actin. Cell 20, 329–341. https://doi.org/10.1016/0092-8674(80)90619-4

29. Mok, L.-P., Qin, T., Bardot, B., LeComte, M., Homayouni, A., Ahimou, F., Wesley, C., 2005. Delta activity independent of its activity as a ligand of Notch. BMC Dev. Biol. 5, 6. https://doi.org/10.1186/1471-213X-5-6

30. Nolo, R., Abbott, L.A., Bellen, H.J., 2000. Senseless, a Zn Finger Transcription Factor, Is Necessary and Sufficient for Sensory Organ Development in Drosophila. Cell 102, 349–362. https://doi.org/10.1016/S0092-8674(00)00040-4

31. Powell, P.A., Wesley, C., Spencer, S., Cagan, R.L., 2001. Scabrous complexes with Notch to mediate boundary formation. Nature 409, 626–630. https://doi.org/10.1038/35054566

32. Renaud, O., Simpson, P., 2001. scabrous modifies epithelial cell adhesion and extends the range of lateral signalling during development of the spaced bristle pattern in Drosophila. Dev. Biol. 240, 361–376. https://doi.org/10.1006/dbio.2001.0482

33. Sakata, T., Sakaguchi, H., Tsuda, L., Higashitani, A., Aigaki, T., Matsuno, K., Hayashi, S., 2004. Drosophila Nedd4 Regulates Endocytosis of Notch and Suppresses Its Ligand-Independent Activation. Curr. Biol. 14, 2228–2236. https://doi.org/10.1016/j.cub.2004.12.028

34. Shaya, O., Binshtok, U., Hersch, M., Rivkin, D., Weinreb, S., Amir-Zilberstein, L., Khamaisi, B., Oppenheim, O., Desai, R.A., Goodyear, R.J., Richardson, G.P., Chen, C.S., Sprinzak, D., 2017. Cell-Cell Contact Area Affects Notch Signaling and Notch-Dependent Patterning. Dev. Cell 40, 505–511.e6. https://doi.org/10.1016/j.devcel.2017.02.009

35. Simpson, P., 1990. Lateral inhibition and the development of the sensory bristles of the adult peripheral nervous system of Drosophila. Dev. Camb. Engl. 109, 509–519. DOI: 10.1242/dev.109.3.509

36. Simpson, P., Woehl, R., Usui, K., 1999. The development and evolution of bristle patterns in Diptera. Dev. Camb. Engl. 126, 1349–1364. DOI: 10.1242/dev.126.7.1349

37. Sokac, A.M., Bement, W.M., 2006. Kiss-and-Coat and Compartment Mixing: Coupling Exocytosis to Signal Generation and Local Actin Assembly. Mol. Biol. Cell 17, 1495–1502. https://doi.org/10.1091/mbc.e05-10-0908

38. Sprinzak, D., Lakhanpal, A., LeBon, L., Santat, L.A., Fontes, M.E., Anderson, G.A., Garcia-Ojalvo, J., Elowitz, M.B., 2010. Cis Interactions between Notch and Delta Generate Mutually Exclusive Signaling States. Nature 465, 86–90. https://doi.org/10.1038/nature08959

39. Troost, T., Schneider, M., Klein, T., 2015. A Re-examination of the Selection of the Sensory Organ Precursor of the Bristle Sensilla of Drosophila melanogaster. PLoS Genet. 11, e1004911. https://doi.org/10.1371/journal.pgen.1004911

